# DJ-1 products glycolic acid and D-lactate restore deficient axonal trafficking and DNA damage response in FUS and SOD1-associated amyotrophic lateral sclerosis

**DOI:** 10.1101/2023.09.20.558653

**Authors:** Arun Pal, Dajana Grossmann, Hannes Glaß, Vitaly Zimyanin, René Günther, Marica Catinozzi, Tobias M. Boeckers, Jared Sterneckert, Erik Storkebaum, Susanne Petri, Florian Wegner, Stefan Grill, Francisco Pan-Montojo, Andreas Hermann

## Abstract

Amyotrophic Lateral Sclerosis (ALS) is the most common motor neuron disease leading to death within 2-5 years. Currently available drugs can only slightly prolong survival. Despite the progress that has been achieved in unravelling the molecular mechanisms of the disease so far, the underlying pathophysiology is not fully understood. We present novel insights into the pathophysiology of *Superoxide Dismutase 1* (SOD1)- and in particular *Fused In Sarcoma* (FUS)-ALS by revealing a putatively central role of the Parkinson’s disease (PD) associated glyoxylase DJ-1 and its products glycolic acid (GA) and D-lactic acid (DL). Combined, but not single, treatment with GA and DL restored axonal trafficking deficits of mitochondria and lysosomes in FUS- and SOD1-ALS patient-derived motoneurons (MNs). This was accompanied by restoration of mitochondrial membrane potential as well as mitochondrial fragmentation (FUS-ALS) or elongation (SOD1-ALS). Furthermore, GA and DL restored cytoplasmic mislocalization of FUS and FUS recruitment to DNA damage sites. We further show that despite presenting an early axonal transport deficiency as well, TDP-43 patient-derived MNs did not share this mechanism. While this points towards the necessity of individualized (gene-) specific therapy stratification, it also suggests common therapeutic targets across different gene variants of ALS. Thus, we introduce a putative novel treatment for ALS based on a combination of the two substances GA and DL which might be not only an interesting novel drug candidate in subsets of ALS cases but also in other neurodegenerative diseases characterized by mitochondrial depolarization.

## Introduction

Amyotrophic Lateral Sclerosis (ALS) is the most common motor neuron disease, with an estimate of 17.000 patients and approximately 5.000 new cases annually only in Europe (Marin, Boumediene et al., 2017). Worldwide incidence is approximately 1.6 cases per 100,000 persons annually (Brotman RG). Compared to other neurodegenerative disorders, ALS exhibits the fastest fatality rate, with an expected survival time of 2–5 years (Brooks, Miller et al., 2000, Brotman RG). Until today, ALS is an incurable disease with Riluzole, Sodium Phenylbutyrate/Taurursodiol and Edaravone being the only approved and commercially available putative disease modifying treatments (Edaravone and Sodium Phenylbutyrate/Taurursodiol not in the EU) (Chen, 2020, Kiernan, Vucic et al., 2021). However, these treatments show limited efficacy (Bensimon, Lacomblez et al., 1994, Chen, Liu et al., 2016, Lacomblez, Bensimon et al., 1996a, Lacomblez, Bensimon et al., 1996b). For example, impacts of Edaravone including real-world settings are mixed, with some studies showing no effects (Witzel, Maier et al., 2022, Writing & Edaravone, 2017) (Abraham, Nefussy et al., 2019, Lunetta, Moglia et al., 2020). Overall, Riluzole can be expected to delay time to death or time to tracheostomy for patients with ALS by about 3 months (Hinchcliffe & Smith, 2017).

ALS appears in familial (∼10% of cases) and sporadic forms (∼90% of cases) (Turner, Al-Chalabi et al., 2017) and is caused by the degeneration of MNs in the spinal cord and brain stem (lower MNs) and the motor cortex (upper MNs), progressively resulting in paralysis and death (Zarei, Carr et al., 2015). Mutations in the genes *C9ORF72, SOD-1, FUS*, and *TARDBP* are the most frequent monogenetic forms associated with familial ALS (Hou, Jiao et al., 2016, Müller, Brenner et al., 2018). The pathogenic mechanisms underlying MN death have been extensively studied. Increased glutamate signaling and intracellular calcium levels (excitotoxicity), ER stress, mitochondrial dysfunction, oxidative stress due to the increase of reactive oxygen species (ROS), dysregulated transcription and RNA processing, protein misfolding and aggregation, dysregulated endosomal trafficking, impaired axonal transport and neuroinflammation are key components involved in the pathogenesis of ALS (for review see (Tadic, Prell et al., 2014, Weishaupt, Hyman et al., 2016)). Nevertheless, the common putative devastating pathophysiological cascade as a whole remains to be understood. However, a common early sign of degeneration is a dying back of the neurons with early axonal trafficking deficits in many genetic ALS forms (Kreiter, Pal et al., 2018, Naumann, Pal et al., 2018, Pal, Glaß et al., 2018, Pal, Kretner et al., 2021). The particular vulnerability of MNs compared to other neuronal groups is still a matter of debate, however, it might involve both high expression of AMPA receptors that lack the calcium impermeable GluR2 subunit, which makes them more prone to excitotoxicity and imbalances in intracellular Ca^2+^ homeostasis (Williams, Day et al., 1997). Moreover, MNs are known to express low levels of Ca^2+^ -buffering proteins which increases their vulnerability (Ince, Stout et al., 1993). Additionally, MNs strongly rely on optimal mitochondrial function, due to their high metabolic demands, and are therefore more prone to cell death when mitochondrial activity is dysregulated. Overall, the crosstalk between Ca^2+^, the endoplasmic reticulum (ER) and mitochondrial function as well as oxidative stress seems to be crucial in the development of ALS pathology (Tadic et al., 2014).

The protein glyoxylase DJ-1, encoded by the *PARK7* gene, is known as a redox-dependent chaperone with neuroprotective potential. Loss-of function mutations cause early-onset autosomal recessive PD (Hague, Rogaeva et al., 2003). DJ-1 overexpression protects dopaminergic neurons against PD, whereas DJ-1 deficiency leads to profound loss of dopaminergic neurons. DJ-1 has several roles including stabilization of the antioxidant master regulator Nuclear Factor Erythroid 2-Related Factor 2 (NRF2) (Clements, McNally et al., 2006). Furthermore, DJ-1 is integral for maintaining mitochondrial potential, Ca^2+^ homeostasis and ATP production. DJ-1 ablation was shown to reduce ER-mitochondria association and disrupted mitochondrial Ca^2+^ uptake (Liu, Ma et al., 2019). Of note, Loss of DJ-1 did not affect mitochondrial respiration but increased ROS production and mitochondrial permeability transition pore opening (Giaime, Yamaguchi et al., 2012), a phenotype, which we recently identified also in FUS-ALS MNs (Zimyanin, Pielka et al., 2023).

Interestingly, we recently showed mitochondrial depolarization as early events in SOD1- and in particular FUS-ALS patients-derived MNs (Gunther, Pal et al., 2022, Naumann et al., 2018). Most importantly, restoration of mitochondrial inner membrane potential delayed neurodegeneration in FUS-ALS (Naumann et al., 2018). ALS-causing mutations in *FUS* are mainly localized in its nuclear localization sequence (NLS) and thus causing a cytoplasmic mislocalization (Japtok, Lojewski et al., 2015, Szewczyk, Gunther et al., 2023, Szewczyk, Gunther et al., 2021). This is accompanied by a loss of nuclear FUS function including a lack of proper recruitment of FUS to DNA damage sites and DNA damage repair (DDR) (Naumann et al., 2018, Wang, Guo et al., 2018), a mechanism downstream of poly(ADP-ribose) polymerase 1 (PARP1) (Mastrocola, Kim et al., 2013, Rulten, Rotheray et al., 2014). Similar to FUS, DJ-1 is also known as putative oncogene (Clements et al., 2006, Naumann, Peikert et al., 2019).

Having shown mitochondrial depolarization as well as mitochondrial transition pore opening without dominant affection of mitochondrial respiration, we hypothesized that similar pathways are disrupted in FUS- and SOD1-ALS as shown for DJ-1 (Toyoda, Erkut et al., 2014). Glycolic acid (GA) and D-lactate (DL) both occur naturally in the cell as products of DJ-1 (Lee, Song et al., 2012). DJ-1 converts the reactive aldehydes glyoxal and methylglyoxal to GA and DL, respectively (Lee et al., 2012, Thornalley, 2003). GA can support the mitochondrial membrane potential and neuronal survival (Toyoda et al., 2014), improve mitochondrial energy production, thereby increasing the levels of NAD(P)H (Bour, Dening et al., 2021) and can also reduce oxidative stress via a glutathione-mediated pathway (Diez, Traikov et al., 2021). Therefore, we speculated that GA and DL might be a treatment option for MNs differentiated from induced pluripotent stem (iPS) cells derived from fibroblasts of ALS patients with different ALS-causing mutations and compared it to the standard of care treatment (Riluzole).

## RESULTS

### GA and DL restrore axonal trafficking in FUS-ALS mutants

Products of the PD-related glyoxalase DJ-1, namely GA and DL, were reported to support mitochondrial membrane potential and neuronal survival in PD animal models (Toyoda et al., 2014). We have recently shown significant mitochondrial dysfunction in FUS-ALS patient-derived MNs, including severe axonal trafficking deficiency and loss of mitochondrial membrane potential (Naumann et al., 2018, Pal et al., 2018). Particularly the latter was rescued by GA and DL in the PD models (Toyoda et al., 2014). Thus, we investigated whether the treatment of GA and DL together is able to rescue FUS-ALS MN phenotypes as well. To this end, patient-derived spinal MNs (Table 1 for details on mutations and patients) were matured for 21 days in vitro (DIV) in microfluidic chambers (MFCs) – time points at which mutant cells showed axon trafficking and mitochondrial phenotypes (Naumann et al., 2018, Pal et al., 2018) – and then treated for 24 h at both sites (distal and proximal) with GA and DL (each 1mM) and imaged using Mitotracker JC-1 and Lysotracker (Fig. 1). The combination of GA and DL did completely restore axon trafficking phenotypes of mitochondria and lysosomes of FUS-ALS spinal MNs (Fig. 1A-C, Fig. S1 for individual cell lines). This was not the case when GA or DL were given individually (Fig. 1A, Fig. S2). While up to 20mM of GA or DL alone had no effect on axon trafficking, the EC_50_ of the combination of GA and DL was 485µM each for restoration of axonal trafficking (Fig. S2). Furthermore, the combinatorial treatment of 1mM GA and DL restored mitochondrial fragmentation (Fig. 1D) and mitochondrial potential (Fig. 1E).

**Figure 1.**
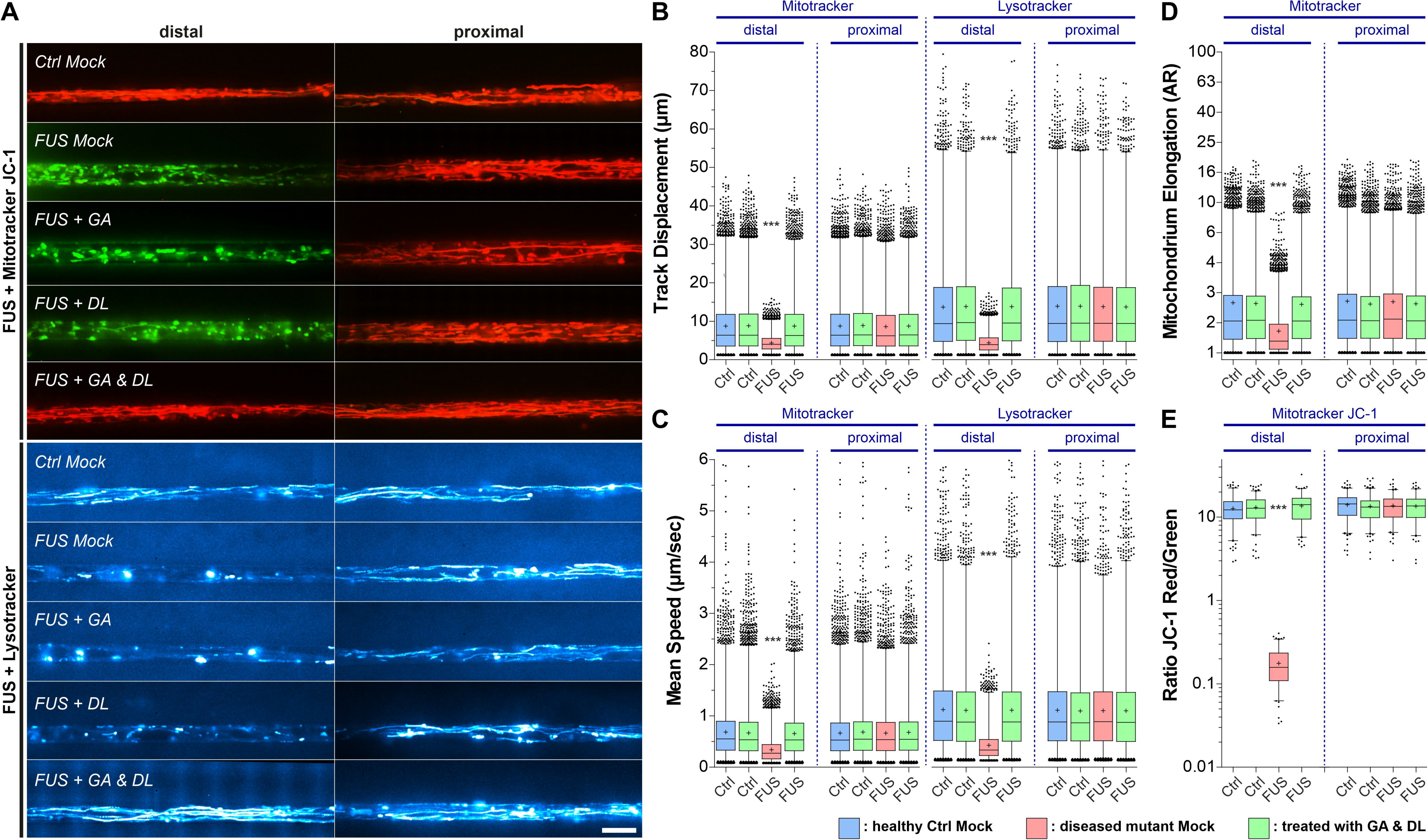
GA and DL together rescue axonal trafficking defects in ALS-FUS mutants. Patient-derived spinal MNs were matured for 21 DIV in MFCs, double treated for 24 h at both sites (distal and proximal) with GA and DL (each 1 mM) and imaged live at the distal (left) vs proximal (right) channel end with Mitotracker JC-1 (red/green) and Lysotracker (cyan hot). **(A)** Maximum intensity projections of movies visualize organelle moving tracks in axons. Processive motility results in straight, longer trajectories whereas immobile organelles project as punctae. Representative examples from the mutant FUS (Fig. S1, Table 1) and control (Ctrl) line pools are shown as follows: FUS: FUS R521C, Ctrl: Ctrl1. Note the exclusively distal loss of lysosomal and mitochondrial motility and its inner membrane potential (JC-1 green) in FUS Mock compared to Ctrl (JC-1 red) which were both rescued through GA and DL double treatment but not through GA or DL alone even at 20 mM (supplementary movies 1, 2). Scale bar = 10 µm. **(B-E)** Box plots quantifications of various tracking and morphology parameters deduced from movies from (A) as per organelle values for the mutant FUS and Ctrl cell line pool, except of (E) showing mean values per image. For mutant FUS, data from the FUS R521C, R521L, R495X and FUS P525L-eGFP lines were pooled (Table 1). For wild-type Ctrl, data from the Ctrl1, Ctrl2, Ctrl3, FUS WT-GFP and SOD1 D90A igc lines were pooled (Table 1). For individual cell lines refer to Fig. S1. Box: 25-75% interquartile range, horizontal line: median, cross: mean, whiskers: non-outlier range (99% of data), dots outside whiskers: outliers, Ctrl Mock is shown in pale blue, diseased mutant mock in pale red, double treatment with GA and DL in pale green. A one-way ANOVA with Bonferroni post-hoc test was utilized to reveal significant differences in pairwise comparisons. Asterisks: highly significant alteration compared to Ctrl Mock distal (in pale blue), *** p ≤ 0.001, all other pairwise comparisons were not significantly different. **(B, C)** Note the drastic reduction in exclusively distal track displacement (B) and mean speed (C) in FUS Mock that was rescued through GA and DL double treatment. **(D)** Note the drastic reduction of exclusively distal mitochondria elongation (fragmentation) in FUS that was rescued by GA and DL double treatment. **(E)** Note the loss of exclusively distal mitochondrial inner membrane potential in FUS that was rescued by GA and DL double treatment.

**Table 1:** Patient/proband characteristics.

| Genotype | Cell line | Sex | Age at biopsy | Mutation | Primarily characterized in |
| --- | --- | --- | --- | --- | --- |
| Wt | Ctrl1 | male | 48 | - | (Japtok et al., 2015) |
| Wt | Ctrl2 | female | 43 | - | (Reinhardt et al., 2013) |
| Wt | Ctrl3 | female | 48 | - | (Reinhardt et al., 2013) |
| IGC | FUS-WT eGFP<sup>het</sup> | isogenic to FUS R521C and FUS-P525L GFP | N/A | - | (Naumann et al., 2018) |
| IGC | SOD1 D90A igc | isogenic to SOD1 D90A | N/A | - | (Bursch et al., 2019) |
| Mt | TDP43 S393L<sup>het</sup> | female | 87 | S393L | (Kreiter et al., 2018) |
| Mt | TDP43 G294V<sup>het</sup> | male | 46 | G294V | (Kreiter et al., 2018) |
| Mt | SOD1 D90A<sup>hom</sup> | female | 46 | D90A | (Naujock et al., 2016) |
| Mt | SOD1 A4V<sup>het</sup> | female | 73 | A4V | (Gunther et al., 2022) |
| Mt | SOD1 R115G<sup>het</sup> | male | 59 | R115G | (Naujock et al., 2016) |
| Mt | FUS R521C<sup>het</sup> | female | 58 | R521C | (Naumann et al., 2018) |
| Mt | FUS R521L<sup>het</sup> | female | 65 | R521L | (Naumann et al., 2018) |
| Mt | FUS R495X<sup>het</sup> | male | 29 | R495QfsX527 | (Naumann et al., 2018) |
| Mt | FUS-P525L eGFP<sup>het</sup> | isogenic to FUS R521C and FUS-WT GFP | N/A | P525L | (Naumann et al., 2018) |
| Wt | HeLa BAC FUS-eGFP WT | female | N/A | - | (Maharana, Wang et al., 2018, Poser et al., 2008) |
| Mt | HeLa BAC FUS-eGFP P525L | Female | N/A | P525L | (Maharana et al., 2018, Poser et al., 2008) |

### GA and DL restore FUS nuclear cytoplasmic mislocalization and recruitment to nuclear laser-irradiated DNA damage sites

ALS-causing mutations in *FUS* are mainly localized in its NLS and thus causing a cytoplasmic mislocalization, i.e. aggregation (Japtok et al., 2015, Szewczyk et al., 2023, Szewczyk et al., 2021). This is accompanied by a loss of nuclear FUS function including a lack of proper recruitment of FUS to DNA damage sites and DNA damage repair (DDR) (Naumann et al., 2018, Wang et al., 2018). First, we investigated whether treatment with GA and DL influenced FUS nuclear/cytoplasmic distribution. To this end, we used engineered HeLa cells expressing a bacterial artificial chromosome (BAC) of either wild-type or P525L mutant FUS tagged with eGFP at the carboxyl-terminus (Poser, Sarov et al., 2008). Similar to axonal trafficking, individual treatment of the P525L mutant with either GA or DL did not influence the count of cytoplasmic FUS aggregates whereas co-treatment with both led to a full rescue back to the wild-type level (Fig. 2A, B). We next used targeted laser irradiation to induce DNA strand breaks at defined nuclear positions as described previously (Naumann, Pal et al. 2018). While mutant FUS cells showed complete loss of FUS recruitment in Mock-treated conditions, this phenotype was restored in case of combined GA and DL treatment but not by single treatments with GA or DL (Fig. 2C, D). The rescue effect of GA and DL was observed at a similar EC_50_ compared to restoration of axonal trafficking (496.5 µM, Fig. S3B) with a remaining minor delay compared to the wild-type control (Fig. 2D, S3A). We finally used CRISPR/CAS9 gene-edited iPSC-derived spinal MNs (Table 1) either expressing wild-type- or P525L FUS-eGFP and proved that 10 mM co-treatment of GA and DL was also able to restore FUS recruitment to laser-irradiated DNA damage sites in spinal MNs (Fig. 2E, F).

**Figure 2.**
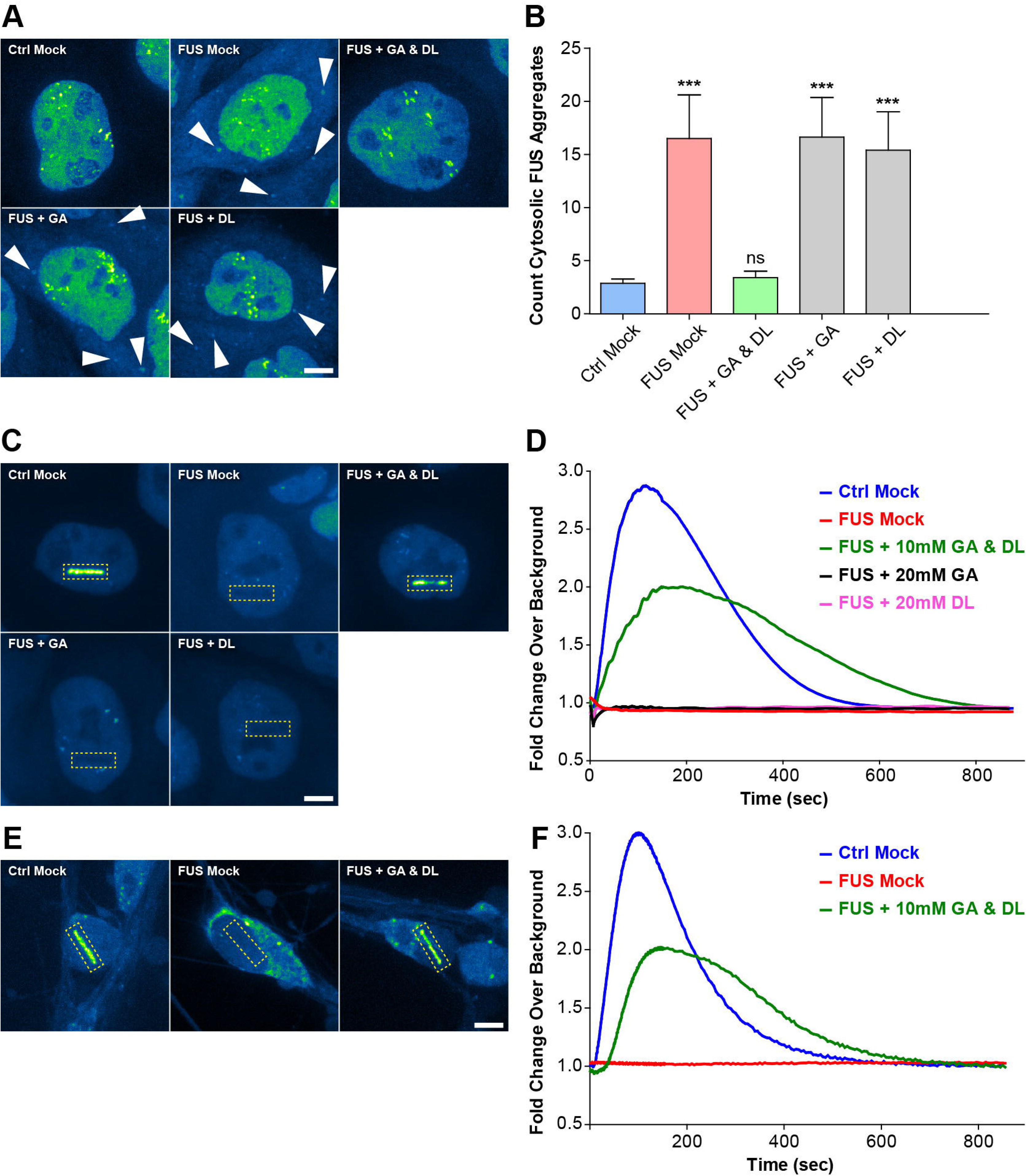
GA and DL together rescue recruitment of FUS to nuclear DNA damage sites in mutant ALS FUS cells. **(A)** GA and DL double treatment rescues from cytosolic FUS aggregation. Shown are maximum intensity projections of confocal Z-stacks imaged live in the transgenic BAC HeLa cell model expressing normal FUS-eGFP WT (Ctrl) or mutant FUS-eGFP P525L (FUS) (Table 1). Note the occurrence of cytosolic FUS aggregates in FUS Mock as compared to Ctrl Mock (arrowheads) that were rescued through GA and DL double treatment but not through GA or DL alone even at 20 mM.Scale bar = 10 µm. **(B)** Quantification of (A) as counts of cytosolic FUS aggregates per cell, N = 30 images, error bars show STDEV. Note the drastic increase in FUS Mock (pale red bar) as compared to Ctrl Mock (pale blue bar) and its reversion back to Ctrl levels through GA and DL double treatment (pale green bar) but not through GA or DL alone (grey bars). A one-way ANOVA with Bonferroni post-hoc test was utilized to reveal significant differences in pairwise comparisons to Ctrl Mock. Asterisks: highly significant alteration in indicated pairwise comparison, *** p ≤ 0.001 and ns: no significant difference. **(C)** Transgenic BAC HeLa cells from (A) were double treated for 24 h with GA and DL (each 10 mM). Recruitment-withdrawal of FUS-GFP to UV Laser cuts in nuclei (boxed area) was then imaged live (supplementary movie 3). Shown are single movie frames at 150 sec when the eGFP intensity was around its maximum. Note the failed recruitment in FUS Mock as compared to Ctrl Mock and its rescue through GA and DL double treatment but not through GA or DL alone even at 20mM. Scale bar = 10µm. **(D)** Quantification of (C). Note the failed recruitment in FUS Mock (red curve) over the entire recording time of 850 sec whereas GA and DL double treatment (green curve) fairly restored the recruitment-withdrawal towards Ctrl kinetics (blue curve) whereas neither GA nor DL alone (black and pink curve, respectively) rescued the FUS-eGFP recruitment. **(E)** Patient-derived isogenic spinal MNs expressing normal (Ctrl) or mutant P525L (FUS) FUS-GFP (Table 1) were matured for 21 DIV and then double treated for 24 h with GA and DL (each 10 mM). Recruitment-withdrawal of FUS-GFP to UV Laser cuts in nuclei (boxed area) was then imaged live (supplementary movie 4). Shown are single movie frames at 150 sec when the eGFP intensity was around its maximum. Note the failed recruitment in FUS Mock as compared to Ctrl Mock and its rescue through GA and DL double treatment. Scale bar = 10 µm. **(F)** Quantification of (E), amount of FUS-eGFP at cut over time. Note the failed recruitment in FUS Mock (red curve) over the entire recording time of 850 sec whereas GA and DL double treatment (green curve) fairly restored the recruitment-withdrawal towards Ctrl kinetics (blue curve).

### Axon trafficking but not FUS DNA damage site recruitment depends primarily on proper mitochondrial function

Both, axon trafficking and DNA damage repair are highly energy demanding and thus depend on the availability of energy in the cell (Martire, Mosca et al., 2015, Naumann et al., 2018). Postmitotic neurons rely mainly on oxidative phosphorylation (OXPHOS) to generate ATP (Zheng, Boyer et al., 2016). We thus interfered with mitochondrial OXPHOS in wild-type control MNs to see whether this is already sufficient to induce axonal trafficking deficiency as well as lack of poly(ADP)ribose-dependent FUS recruitment to DNA damage sites. Wild-type MNs (Ctrl 1-3, Table 1) were treated with 10 µM Oligomycin A, an inhibitor of the respiratory chain complex V, or with 10 µM of the uncoupler carbonyl cyanide 3-chlorophenylhydrazone (CCCP) for 24 hours on the distal site of MFCs. The resulting dysfunction of axonal mitochondria induced phenocopies of FUS-ALS axonal trafficking defects (Fig. 3A). Conversely, the same treatments did not interfere with FUS-recruitment to laser-induced DNA damage sites in MNs expressing wild-type FUS-eGFP (Table 1) (Fig. 3B, C), albeit an increase in cytosolic FUS (Fig. 3D) that was insufficient to interfere with its nuclear function.

**Figure 3.**
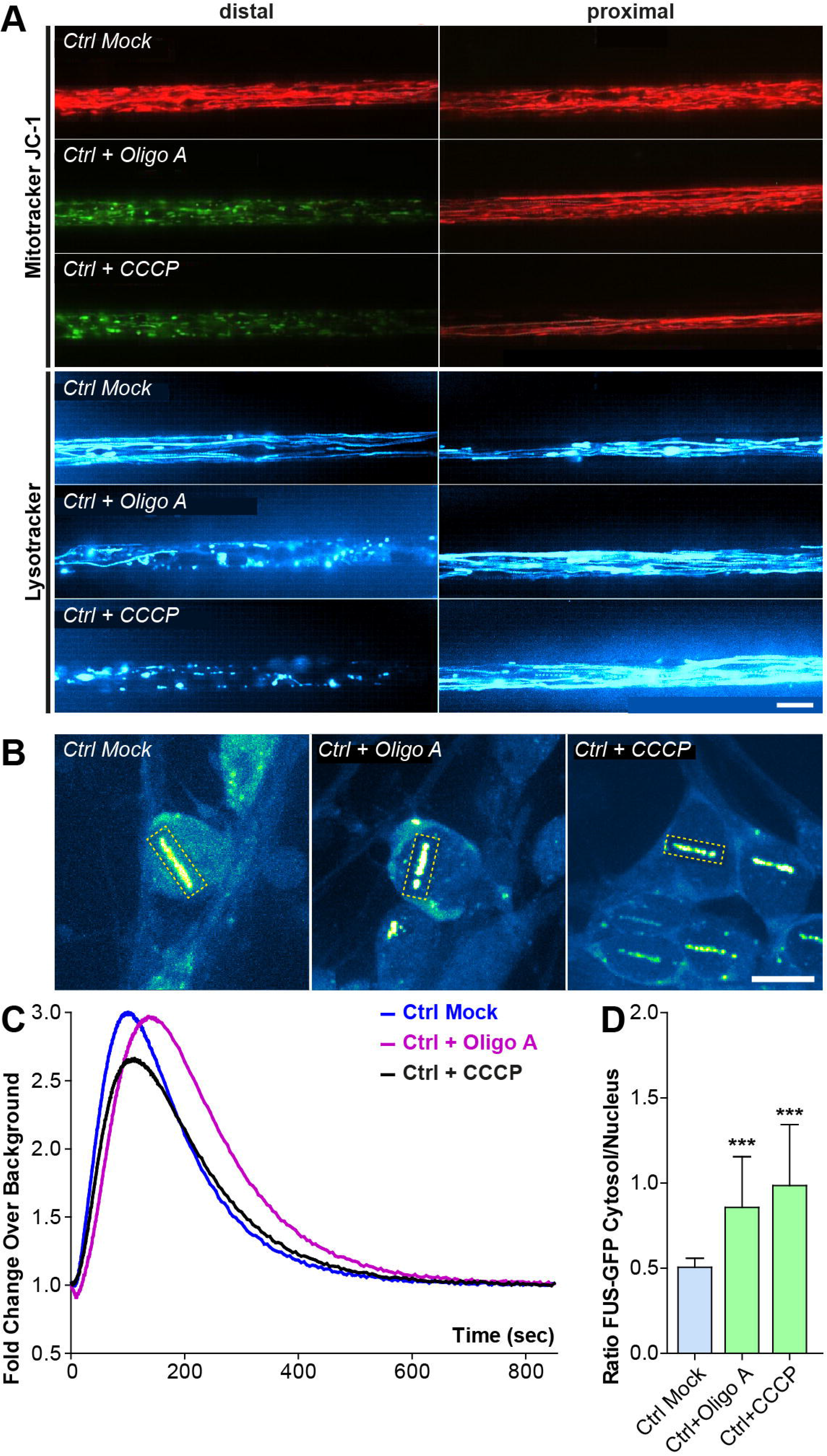
Site-specific inhibition of mitochondrial ATP production causes local disruption of axonal organelle trafficking. **(A)**. Patient-derived spinal MNs from healthy control patients were matured for 21 DIV in MFCs. Then 10µM of Oligo A or CCCP was exclusively added to the distal site 4 h prior to live imaging at the distal (left) vs proximal (right) channel end with Mitotracker JC-1 (red/green) (supplementary movie 5) and Lysotracker (cyan hot) (supplementary movie 6). Shown are maximum intensity projections of movies to visualize organelle moving tracks in axons. Processive motility results in straight, longer trajectories whereas immobile organelles project as punctae. Note the drastic loss of lysosomal and mitochondrial motility and its inner membrane potential (JC-1 green) at the treated distal site (Oligo A, CCCP) compared to Mock whereas the untreated proximal site remained physiological. Shown is a representative example of Ctrl1 (Table 1). Scale bar = 10 µm. **(B)** Patient-derived spinal MNs expressing normal (Ctrl) FUS-GFP WT (Table 1) were matured for 21 DIV in uncompartmentalized dishes. Cells were treated for 4 h with 10µM Oligo A or CCCP, recruitment-withdrawal of FUS-eGFP to UV Laser cuts in nuclei was then imaged live (supplementary movie 7) and **(C)** the quantified amount of FUS-eGFP at cuts plotted over time. Note that neither Oligo A (purple curve) nor CCCP (black curve) inhibited the normal FUS recruitment (blue curve). **(D)** Quantification of (B) of the ratio of FUS-eGFP cytosolic over nuclear integral intensity, N=30 images, error bars show STDEV. A one-way ANOVA with Bonferroni post-hoc test was utilized to reveal significant differences in pairwise comparisons to Ctrl Mock. Asterisks: highly significant alteration in indicated pairwise comparison, *** p ≤ 0.001.

### DJ-1 products GA and DL restore nuclear phenotypes by restoring NAD metabolism

We recently reported that restoration of nuclear phenotypes in FUS-ALS MNs subsequently led to a restoration of axonal trafficking and disappearance of cytosolic FUS aggregates (Naumann et al., 2018, Pal et al., 2018). Therefore, we were wondering about the mechanisms by which GA and DL restore nuclear functions in FUS-ALS including proper recruitment of FUS to sites of DNA damage. FUS is recruited to DNA damage sites downstream of PARP1 (Mastrocola et al., 2013, Rulten et al., 2014) which is impaired in case of ALS-causing mutations in the *FUS* gene (see Fig. 2, S3 and (Naumann et al., 2018)). Poly(ADP-ribose) polymerases belong to the three main enzymes consuming NAD^+^. Mammalian cells have evolved a NAD^+^ salvage pathway capable to resynthesize NAD^+^. The products of the glyoxylase DJ-1 – GA and DL – can be reintroduced into metabolic pathways. D-lactic acid can be converted to pyruvate reducing NAD(P)^+^ to NAD(P)H via the D-Lactate dehydrogenase, while GA converted to glyoxylate (for further use in the citrate cycle) also reduces NAD(P)^+^ to NAD(P)H. Therefore, we hypothesized that supplementation with the DJ-1 products GA and DL restores NAD^+^ using this salvage pathway and therefore the NAD^+^ precursor nicotinamide riboside (NAR) (Harlan, Killoy et al., 2020, Harlan, Pehar et al., 2016) leads to a similar rescue effect as GA and DL.

To this end, we supplemented the BAC HeLa FUS model (Table 1) either with NAR or inhibited the rate limiting enzyme of NAD(P)^+^ synthesis, nicotinamide phosphoribosyltransferase (NAMPT), using FK866. NAR supplementation rescued FUS cytoplasmic mislocalization (Fig. 4A, B) and also FUS recruitment to DNA damage sites in the FUS-eGFP P525L mutant (Fig. 4C) in a dose-dependent manner (Fig. 4E). Furthermore, NAR increased the abundance of FUS at DNA damage sites in wild-type cells (Fig. 4D). Conversely, inhibition of NAMPT with FK866 led to a FUS-ALS phenocopy with cytoplasmic mislocalization of FUS (Fig. 4A, B) and dose-dependent loss of recruitment of FUS to sites of DNA damages in wild-type cells (Fig. 4C, F) while it had no additional effect on mutant FUS cells (Fig. 4C, G).

**Figure 4.**
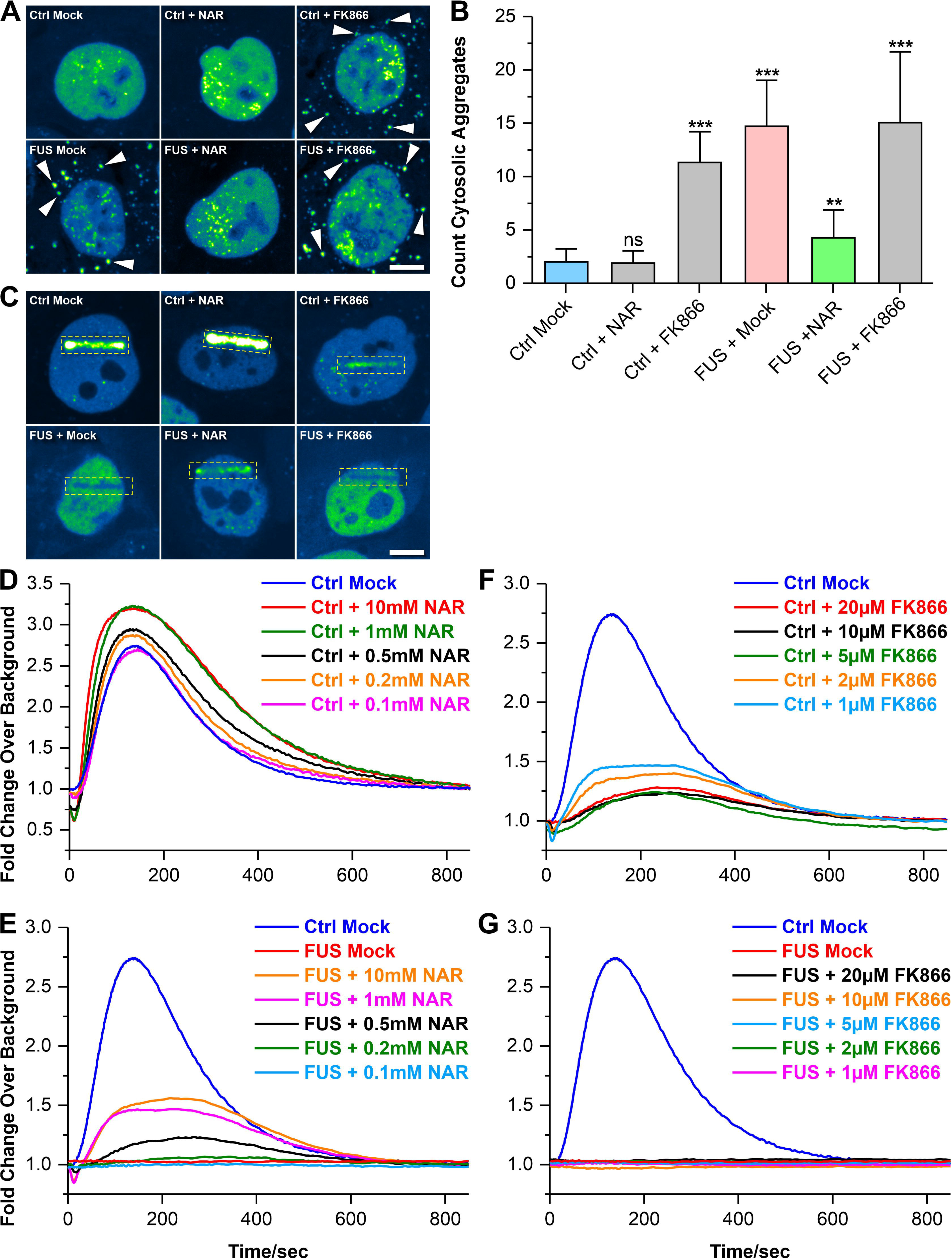
Recruitment of FUS to nuclear DNA damage sites depends on proper NAD metabolism. **(A)** Shown are maximum intensity projections of confocal Z-stacks imaged live in the transgenic BAC HeLa cell model expressing normal FUS-eGFP WT (Ctrl) or mutant FUS-eGFP P525L (FUS) (Table 1). Note the occurrence of cytosolic FUS aggregates in FUS Mock as compared to Ctrl Mock (arrowheads) that were rescued through treatment with 10mM nicotinamide riboside (NAR) over 24 h, a precursor of NAD^+^. Conversely, inhibition of NAD synthesis through treatment with 10µM FK688 over 24 h phenocopied the cytosolic mutant FUS aggregation. Scale bar = 10 µm. **(B)** Quantification of (A) as counts of cytosolic FUS aggregates per cell, N = 30 images, error bars show STDEV. Note the drastic increase in FUS Mock (pale red bar) as compared to Ctrl Mock (pale blue bar) and its reversion back to Ctrl levels through NAR treatment (pale green bar) whereas FK866 treatment of Ctrl cells led to increased cytosolic FUS aggregation similar to mutant FUS Mock. A one-way ANOVA with Bonferroni post-hoc test was utilized to reveal significant differences in pairwise comparisons to Ctrl Mock. Asterisks: highly significant alteration in indicated pairwise comparison, ** p ≤ 0.01, *** p ≤ 0.001 and ns: no significant difference. **(C)** Transgenic BAC HeLa cells from (A) were treated for 24 h with either NAR or FK866. Recruitment-withdrawal of FUS-GFP to UV Laser cuts in nuclei (boxed area) was then imaged live (supplementary movie 8). Shown are single movie frames at 150 sec when the GFP intensity was around its maximum. Note the failed recruitment in FUS Mock as compared to Ctrl Mock and its rescue through NAR treatment whereas FK866 treatment of Ctrl cells decreased the FUS recruitment. Scale bar = 10 µm. **(D)** Quantification of (C) for Ctrl cells, GFP intensities at cuts plotted over time after Laser irradiation. Note the further boosted recruitment of FUS over Ctrl Mock (blue curve) through NAR treatment in a concentration-dependent manner. **(E)** Quantification of (C) for mutant FUS cells. Note the rescue of FUS recruitment over Mock (red curve) through NAR treatment in a concentration-dependent manner, even though not fully restored to Ctrl Mock levels (blue curve). **(F)** Quantification of (C) for Ctrl cells. Note the pronounced decrease of FUS recruitment through FK866 treatment in a concentration-dependent manner, even though no complete inhibition as in FUS Mock was achieved. **(G)** Quantification of (C) for mutant FUS cells. Note that FK866 treatment did not alter the failed FUS recruitment at any concentration.

### GA and DL restore axonal trafficking in SOD1-but not TDP43-ALS mutants

We finally wished to test whether the potential therapeutic effects of GA & DL are also seen in other monogenetic ALS mutants. To this end, we chose SOD1- and TDP43-mutant iPSC-derived spinal MNs as these have been reported to also show axonal trafficking phenotypes at DIV 21, albeit clearly distinct ones (Gunther et al., 2022, Kreiter et al., 2018, Pal et al., 2018): While mutant SOD1 MNs exhibited an elongation of mitochondria along with a reduction of the inner membrane potential but no alteration of mitochondrial and lysosomal speed and track displacement (Gunther et al., 2022), mutant TDP43 MNs displayed a striking decrease in speed and track displacement of both types of organelles in distal and proximal axons but no perturbed mitochondrial inner membrane potential (Kreiter et al. 2018, Pal et al. 2018). Treatment with 1mM GA and DL for 24 hours was able to restore the axonal mitochondrial phenotypes of SOD1 mutant patient-derived spinal MNs in both the distal and proximal axon parts (Fig. 5A, D, E, S1) but not those of mutant TDP43 spinal MNs (Fig. 5A-C, S1). Specifically, adding 1mM GA and DL restored the mitochondrial elongation and depolarization in SOD1 mutant MNs (Fig. 5A, D, E). Thus, responsiveness to GA and DL seems to be associated with mitochondrial depolarization, as shown for our FUS (Fig. 1A, E, S1C) and SOD1 (Fig. 5A, E, S1C) mutants.

**Figure 5.**
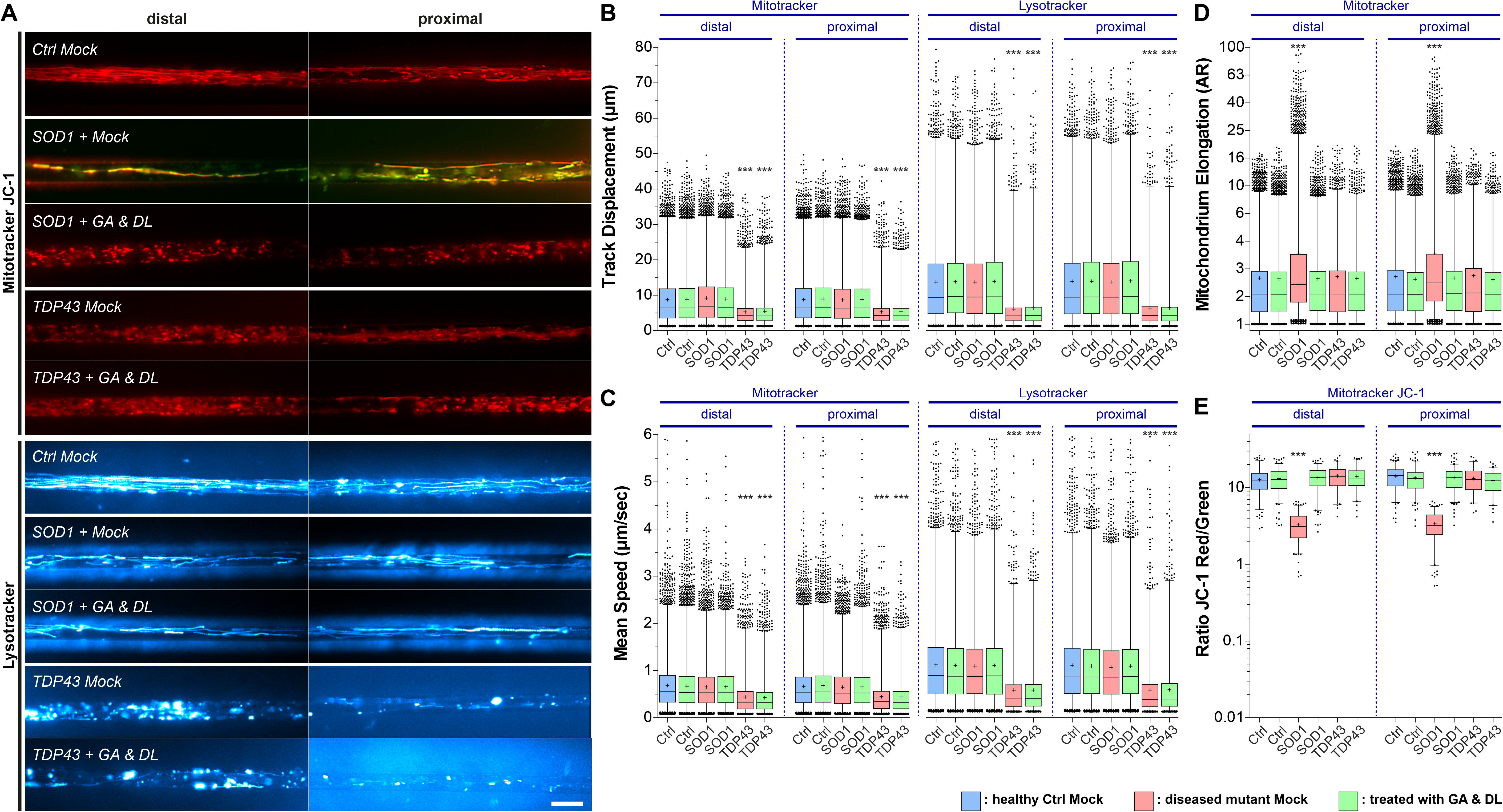
GA and DL together rescue axonal organelle phenotypes in ALS-SOD1 but not in TDP-43 mutants. Patient-derived spinal MNs were matured for 21 DIV in MFCs, double-treated for 24 h at both sites (distal and proximal) with GA and DL (each 1 mM) and imaged live at the distal (left) vs proximal (right) channel end with Mitotracker JC-1 (red/green) and Lysotracker (cyan hot). **(A)** Maximum intensity projections of movies visualize organelle moving tracks in axons. Processive motility results in straight, longer trajectories whereas immobile organelles project as punctae. For mitochondria in SOD1, the first frame of the movie is shown instead of the maximum intensity projection to better document the elongation of these organelles. Representative examples from the mutant SOD1 and TDP-43 (Fig. S1, Table 1) and control (Ctrl) line pools are shown as follows: SOD1: SOD1 D90A, TDP-43: TDP-43 S393L, Ctrl: Ctrl1. Note the striking elongation of mitochondria along with a reduction of the inner membrane potential (JC-1 yellow overlap) in SOD1 Mock compared to Ctrl (JC-1 red) at both the distal and proximal site which were rescued through GA and DL double treatment. Conversely, mitochondrial inner membrane potential appeared normal in TDP-43 Mock (JC-1 red) whereas mitochondrial and lysosomal motility was reduced at both the distal and proximal site (punctae instead of trajectories) and could not be restored through GA and DL double treatment (supplementary movies 9, 10). Scale bar = 10 µm. **(B-E)** Box plots quantifications of various tracking and morphology parameters deduced from movies from (A) as per organelle values for the mutant SOD1, TDP-43 and Ctrl cell line pool, except of (E) showing mean values per image. For mutant SOD1, data from the SOD1 D90A, A4V and R115G lines were pooled (Table 1). For mutant TDP-43, data from the TDP-43 S393L and G294V lines were pooled (Table 1). For wild-type Ctrl, data from the Ctrl1, Ctrl2, Ctrl3, FUS WT-GFP and SOD1 D90A igc lines were pooled (Table 1). For individual cell lines refer to Fig. S1. Box: 25-75% interquartile range, horizontal line: median, cross: mean, whiskers: non-outlier range (99% of data), dots outside whiskers: outliers, Ctrl Mock is shown in pale blue, diseased mutant mock in pale red, double treatment with GA and DL in pale green. A one-way ANOVA with Bonferroni post-hoc test was utilized to reveal significant differences in pairwise comparisons. Asterisks: highly significant alteration compared to Ctrl Mock distal (in pale blue), *** p ≤ 0.001, all other pairwise comparisons were not significantly different. **(B, C)** Note the drastic reduction in distal as well as proximal track displacement (B) and mean speed (C) in TDP-43 Mock that was not rescued through GA and DL double treatment. **(D)** Note the drastic elongation of mitochondria at both the distal and proximal site in SOD1 that was rescued by GA and DL double treatment. **(E)** Note the reduction of mitochondrial inner membrane potential in SOD1 at both the distal and proximal site that was rescued by GA and DL double treatment.

### Riluzole did not restore mitochondrial phenotypes in FUS- and SOD1- ALS mutant MNs

We finally investigated whether the “gold standard” treatment of ALS, Riluzole, has similar beneficial effects on axonal trafficking in FUS- and SOD1-patient-derived MNs. To this end, we treated FUS- and SOD1- mutant iPSC-derived spinal MNs with 10 µM Riluzole over the entire maturation period in MFC’s until 21 DIV prior imaging. Interestingly, Riluzole had no effect on any of the parameters investigated, which included mitochondrial and lysosomal trafficking (Fig. S4A-C), mitochondrial inner membrane potential (Fig. S4E) and mitochondrial fragmentation (FUS-ALS) or elongation (SOD1-ALS) (Fig. S4D).

### Discussion

In this study, we introduce a putative novel treatment for ALS based on a combination of the two substances glycolic acid (GA) and D-lactate (DL). Both compounds occur naturally in the cell as products of DJ-1 (Lee et al., 2012), which converts the reactive aldehydes glyoxal and methylglyoxal to GA and DL, respectively (Lee et al., 2012, Thornalley, 2003). We show that combinatorial treatment with GA and DL restored axonal trafficking deficits of mitochondria and lysosomes in FUS- and SOD1-ALS. This was accompanied by restoration of mitochondrial membrane potential as well as mitochondrial fragmentation in case of FUS-ALS or elongation in case of SOD1-ALS. GA and DL furthermore restored cytoplasmic mislocalization of FUS and FUS recruitment to DNA damage sites. Of note, these effects were only seen in case of mitochondrial depolarization (i.e. FUS-ALS and SOD1-ALS) but not in TDP43-ALS, in which mitochondrial membrane potential is not disturbed. This fits to previous data on *PARK7* cells and C. elegans models, in which GA restored the mitochondrial membrane potential and prolonged neuronal survival (Toyoda et al., 2014). It remains, however, open, why in case of ALS mutants, only a combinatorial treatment of GA and DL was able to restore the phenotypes.

GA and DL can help to maintain calcium homeostasis (Chovsepian, Berchtold et al., 2022). ER-mitochondria associations have become of increasing interest in neurodegenerative diseases, since these specialized tight structural associations between a closely apposed ER surface and outer mitochondria membrane were reported to regulate a variety of essential physiological functions including calcium signaling, phospholipid synthesis/exchange and mitochondrial biogenesis and dynamics and cell death (Grossmann, Berenguer-Escuder et al., 2019, Grossmann, Malburg et al., 2023, Liu et al., 2019, Pereira, Grossmann et al., 2023). Interestingly, loss of DJ-1 led to reduced ER-mitochondria association and disturbed function of mitochondria-associated membranes (MAM) and mitochondria in vitro (Liu et al., 2019). Whether the beneficial effects of GA and DL in FUS- and SOD1-ALS are attributable to improved mitochondria-ER interactions requires further investigations.

Apart from rescuing mitochondrial depolarization, fragmentation/elongation and axonal transport, GA and DL did also restore FUS recruitment to laser induced DNA damage sites (Fig. 2). We recently showed that the lack of proper FUS-recruitment to DNA damage sites is upstream of all axonal/mitochondrial phenotypes in FUS-ALS (Naumann et al., 2018). DNA damage induces PARP1, an enzyme important in initiating proper DNA damage response. Activation of PARP1, however, not only leads to NAD^+^ depletion but also to mitochondrial dysfunction (Szabo, Zingarelli et al., 1996). Silencing of PARP1 increased basal cellular parameters of OXPHOS, providing direct evidence that PARP1 is a regulator of mitochondrial function in resting cells. While PARP1 is a regulator of OXPHOS in resting and oxidatively stressed cells, it only exerts a minor effect on glycolysis (Modis, Gero et al., 2012). Interestingly though, energy depletion itself is not sufficient to induce DNA damage (Szabo et al., 1996). This perfectly fits to our data showing that administration of Oligomycin A or CCCP led to mitochondrial depolarization and halted axonal transport whereas it did not influence laser-induced FUS-recruitment (Fig. 3B, C). Collectively, we suggest that the lack of proper DNA repair in FUS-ALS leads to a sustained PARP1 activation and NAD^+^ degradation.

Glyoxylases are important to detoxify e.g. methylglyoxal (MGO) and glyoxal (GO), which are generated e.g. during glycolysis and which are further augmented by ROS. If not detoxified, advanced glycation end products (AGEs) are accumulating. Inceased levels of AGEs are associated with aging and with diverse disorders such as diabetes, renal failure and neurodegeneration. Lipid peroxidation and AGEs occur in the brain during normal aging as well as in Alzheimer’s disease (Dei, Takeda et al., 2002) and also in ALS patient spinal cord motor neurons (Kikuchi, Ogata et al., 2000, Kikuchi, Shinpo et al., 2002). AGE Nε-(carboxymethyl) lysine levels are elevated in cerebrospinal fluid of ALS patients (Kaufmann, Boehm et al., 2004).

The generation of MGO and GO is further augmented by ROS. The transcription factor NRF2 is a critical inducer of the antioxidant response element (ARE)-mediated gene expression, and, importantly, is regulated by DJ-1. Overexpression of DJ-1 results in increased NRF2 protein levels by preventing association with its inhibitor protein, KEAP1, and subsequent ubiquitination of NRF2 (Clements et al., 2006). It is of note that increased NRF2 levels protect against DNA damage by activating DNA damage repair factors. Interestingly, previous data of SOD1 mice showed that the toxicity of astrocytes expressing ALS-linked mutant hSOD1 to co-cultured motor neurons was reversed by NRF2 overexpression (Vargas, Johnson et al., 2008). Since GA and DL are both products of the glyoxylase DJ-1 and most likely act downstream of DJ-1 and NRF2 or independent of the latter, we wondered whether effects of GA and DL were due to alleviation of impaired metabolism by reintroducing them into metabolic pathways. DL can be converted to pyruvate reducing NAD(P)^+^ to NAD(P)H, while GA converted to glyoxylate (for further use in the citrate cycle) also reduces NAD(P)^+^ to NAD(P)H. NAD^+^ is an important metabolite in human cells pivotal for processes including DNA repair and mitophagy (Hou, Wei et al., 2021). Interestingly, enzymes involved in NAD^+^ salvage, namely NAMPT and NMNAT2, were reported to show an altered expression in the spinal cord of ALS patients, suggesting deficits of this pathway in the human ALS pathology (Harlan et al., 2020). Therefore, we hypothesized that supplementation with the DJ-1 products GA and DL restores NAD^+^ using this salvage pathway and thus, boosting this salvage pathway with the NAD^+^ precursor NAR will have similar effects as GA and DL treatment (Harlan et al., 2020, Harlan et al., 2016). Fitting to this theory, inhibition of NAMPT perfectly phenocopied FUS-ALS mutants whereas supplementation with NAR rescued FUS-ALS phenotypes (Fig. 4). Of note, enhancing the NAD^+^ salvage pathway was recently reported to revert the toxicity of astrocytes expressing ALS-linked mutant hSOD1 to co-cultured MNs (Harlan et al., 2016). Conversely, knock out of DJ-1 in the G93A SOD1 ALS mouse model led to an accelerated disease course and shortened survival (Lev, Barhum et al., 2015). In addition, it has been recently shown that increased demand for NAD^+^ relative to ATP induces aerobic glycolysis (Luengo, Li et al., 2021). We recently showed that a boosted metabolic turnover of the glycolytic pathway improved the viability of FUS-ALS MN, while blocking glycolysis reduced their viability (Zimyanin et al., 2023). These data alltogether suggest a metabolic rescue effect through GA and DL treatment in FUS- and SOD1-ALS MNs.

The natural compound d-lactate is found in the body at concentrations of about of 10-20 µM in the blood serum (Hasegawa, Fukushima et al., 2003). Physiological serum concentrations of glycolate are around up to 12.5 µM and a bit higher in tissues (also in brain tissue)(Hagen, Walker et al., 1993, Knight, Hinsdale et al., 2012). The concentrations found e.g. in diabetic patients are even higher, however GA and DL are thought to represent the products of detoxified MGO and GO, respectively, and not the toxic agents themselves. Thus, the respective EC50 found in our study were considerably higher and thus future evaluation of their safety are needed.

In summary, we present novel insights into the pathophysiology of SOD1- and particularly FUS-ALS revealing a putatively central role of the PD-associated glyoxylase DJ-1 and its products GA and DL. We also show that albeit presenting an early axonal transport deficiency as well, TDP-43 patient-derived MNs did not share this mechanism. This spoint towards the necessity of individualized (gene-) specific therapy stratification. This finding also suggests that mitochondrial depolaraization (found e.g. in FUS, SOD1-ALS, but also DJ-1-PD and others) might be a common drug target. GA and DL might thus constitute interesting novel drug candidates in subsets of ALS cases and feasibly other neurodegenerative diseases suffering from mitochondrial depolarization.

## Materials and Methods

### Characteristics of patients for iPSC derivation

We studied iPSC-derived spinal MN cell cultures from familiar ALS patients with the following pathogenic mutations (Mt): TDP43 S393L^het^, TDP43 G294V^het^, SOD1 D90A^hom^, SOD1 A4V^het^, SOD1 R115G^het^, FUS R521C^het^, FUS R521L^het^, FUS R495QfsX527^het^ and compared them to MNs carrying human wildtype counterpart alleles in three cell lines from healthy volunteers (WT, Ctrl1-3), and a gene-corrected isogenic control (IGC) line of SOD1 D90A (SOD1 D90A igc). Moreover, parental FUS R521C was used to generate isogenic FUS-P525L eGFP and its gene-corrected control FUS-WT eGFP(Naumann et al., 2018). All cell lines were obtained by skin biopsies of patients and healthy volunteers and have been described before (Bursch, Kalmbach et al., 2019, Japtok et al., 2015, Kreiter et al., 2018, Naujock, Stanslowsky et al., 2016, Naumann et al., 2018) (Table 1). The performed procedures were in accordance with the Declaration of Helsinki (WMA, 1964) and approved by the Ethical Committee of the Technische Universität Dresden, Germany (EK 393122012 and EK 45022009). Written informed consent was obtained from all participants for publication of any research results.

### Genotyping

DNA from the cell lines were genotyped by a diagnostic human genetic laboratory (CEGAT, Tübingen, Germany). Control lines were also genotyped and did not show any ALS-associated mutation.

### Mycoplasma testing

Every cell line was checked for mycoplasma when entering the lab and after reprogramming., Routine checks for mycoplasma were done every three to six months. We used the Mycoplasma Detection kit according to manufacturer’s instructions (Firma Venor GeM, No 11–1025).

### Generation, gene-editing and differentiation of human iPSC cell lines to MNs in microfluidic chambers (MFCs)

The generation and expansion of iPSC lines from healthy control and familiar ALS patients with defined mutations in distinct ALS genes (Table 1) were recently described (Japtok et al., 2015, Naujock et al., 2016, Naumann et al., 2018). The gene-corrected isogenic control to the homozygous mutant SOD1 D90A (SOD1 D90 igc, table 1) was generated by CRISPR/- Cas9-mediated gene-editing and fully characterized (Bursch et al., 2019). To generate the two isogenic cell lines FUS-WT eGFP and FUS-P525L eGFP, the FUS R521C line was used as parental source (Table 1). The patient-specific FUS R521C mutation was altered at its mutation site and simultaneously C-terminally tagged with eGFP by CRISPR/Cas9-mediated genome editing to obtain a P525L mutation instead along with a gene-corrected wild-type control. Both new lines were fully characterized (Naumann et al., 2018). The subsequent differentiation of all iPSC lines to neuronal progenitor cells (NPC) and further maturation to spinal MNs was described previously (Naumann et al., 2018, Reinhardt, Glatza et al., 2013). The coating and assembly of MFCs (Xona Microfluidics RD900) to prepare for the seeding of MNs was performed as described (Naumann et al., 2018, Pal et al., 2018, Pal et al., 2021). MNs were seeded for maturation into one site of a MFC to obtain a fully compartmentalized culture with proximal somata and their dendrites being physically separated from their distal axons as only the latter type of neurite was capable to grow from the proximal seeding site through a microgroove barrier of 900 µm-long microchannels to the distal site (shown in details in (Glaß, Neumann et al., 2020)). All subsequent imaging in MFCs was performed at DIV 21 of axon growth and MN maturation (DIV 0 = day of seeding into MFCs).

### Live imaging of MNs in MFCs

Time-lapse movie acquisition was performed as described previously (Naumann et al., 2018, Pal et al., 2018). In brief, to track lysosomes and mitochondria, cells were double-stained with live cell dyes Lysotracker Red DND-99 (Molecular Probes Cat. No. L-7528) and Mitotracker Deep Red FM (Molecular Probes Cat. No. M22426) at final concentrations of 50nM each. Trackers were added from a 1 mM stock in DMSO directly to culture supernatants and incubated for 1 h at 37°C. Live imaging was then performed without further washing of cells in the Center for Molecular and Cellular Bioengineering, Technische Universität Dresden (CMCB) light microscopy facility on a Leica HC PL APO 100x 1.46 oil immersion objective on an inversed fluorescent Leica DMI6000 microscope enclosed in an incubator chamber (37°C, 5% CO2, humidified air) and fitted with a 12-bit Andor iXON 897 EMCCD camera (512x512 pixel, 16 µm/pixels on chip, 229.55 nm/pixel at 100x magnification with intermediate 0.7X demagnification in the optical path through the C-mount adapter connecting the camera with the microscope). For more details, refer to https://www.biodip.de/wiki/Bioz06_-_Leica_AFLX6000_TIRF and our previous publications (Naumann et al., 2018, Pal et al., 2018). Fast dual color movies were recorded at 3.3 frames per second (fps) per channel over 2 min (400 frames per channel in total) with 115 ms exposure time as follows: Lysotracker Red (excitation: 561nm Laser line, emission filter TRITC 605/65 nm) and Mitotracker Deep Red (excitation: 633nm Laser line, emission filter Cy5 720/60 nm). Dual channel imaging was achieved sequentially by fast switching between both laser lines and emission filters using a motorized filter wheel to eliminate any crosstalk between both trackers.

### Organelle tracking and shape analysis of live imaging movies

Recently, we have published a comprehensive description of the automated analytical pipeline starting from object recognition in raw movie data to final parametrization of organelle motility and morphology (Pal et al., 2018). In brief, organelle recognition and tracking was performed with the FIJI Track Mate plugin, organelle shape analysis with our custom-tailored FIJI Morphology macro that is based on the FIJI particle analyzer. Both Track Mate and particle analyzer tools returned the mean speed and track displacement for each organelle type (Mito- versus Lysotracker-labeled) along with the length of mitochondria (elongation). Subsequent data mining of individual per-movie result files was performed in KNIME to assemble final results files with annotated per-organelle parameters, thereby allowing all data from each experimental condition to be pooled (e.g. all data for mitotracker or lysotracker for a given cell line). Data per-organelle were visualized as box plots (Fig. 1B- D, 5B-D, S1A, B, D, S2A) except of Fig. 1E, 5E and S1C (ratio red/green Mitotracker JC-1, see below) that show averaged per-movie data. Data for all box plots were pooled from four independent experiments.

### Analysis of inner mitochondrial membrane potential (ratio JC-1 red/green channel)

Analysis of mitochondrial membrane potential with Mitotracker JC-1 (Molecular Probes Cat. No. M34152) was performed as described previously (Naumann et al., 2018). In brief, object segmentation was performed with the channel of higher intensity (most often red emission) to generate a selection limited to mitochondria using a custom-tailored FIJI macro. The resulting selection was saved as a region of interest (ROI) and applied to both channels to reveal the total integral intensity and area of mitochondria and background in both channels using the “Measure” command. After area normalization and background subtraction, ratios of integral red/green intensity were taken as mean membrane potential per movie (first frame only) and batch-analyzed in KNIME as for the tracking analysis (see above). The resultant ratios were displayed as box plots on a log scale (Fig. 1E, 5E, S1C).

### Image/movie quantification

For fluorescent FUS-eGFP aggregation (Fig. 2B, 4B) in HeLa cells, three independent experiments were performed and 10 confocal Z-stacks per experiments analyzed as described previously (Naumann et al., 2018). For movie analysis of MFCs (organelle tracking and shape, mitochondrial inner membrane potential), at least ten movies were acquired of each MFC (= one technical replicate) with three MFCs per experiment and four independent experiments (= MN differentiation pipeline) per cell line.

### Statistical analyses of box plots and bar graphs

Statistical analyses were performed using GraphPad Prism version 5.01. To test for significance between multiple groups, one-way ANOVA followed by Bonferroni’s multiple comparisons test was used. Alpha < 0.05 was used as the cut off for significance (*p < 0.05, **p < 0.005, ***p < 0.001, ****p < 0.0001). For bar graphs, all values are given as mean ± SD.

### DNA damage laser irradiation assay

Isogenic FUS-WT eGFP and FUS-P525L eGFP (Table 1) spinal MNs were differentiated from NPCs as described above, finally split and 300,000 cells seeded into uncompartmentalized 3.5 cm dishes instead of MFCs. Dishes were coated before with poly-L-lysine and laminin as described. All subsequent imaging of DNA damage response to laser irradiation sites was performed at DIV 21 of MN maturation (D0 = day of seeding into final dishes) as described previously (Naumann et al., 2018). In brief, a focused 355 nm UV laser beam was directed through a stereotactic galvanometric mirror box to desired x-y-z-Positions in cell samples held on a standard inverted Axio Observer Z1 Zeiss microscope equipped with a motorized stage and a piezo-electric Z-actuator. A Zeiss alpha Plan-Fluar 100 × 1.45 oil immersion objective was used and 24 laser shots in 0.5μm-steps were administered over 12μm linear cuts located within cell nuclei. The cellular response to this DNA damage comprised a fast recruitment of FUS-eGFP to the laser cut site followed by its slower withdrawal (on-off kinetics) and were recorded live over 15 min by confocal spinning disc imaging of the eGFP tag using a 488 nm laser line and a 12-bit Andor iXON 897 EMCCD camera (512×512, 16 μm pixels, 229.55 nm/pixel at 100Χ magnification) at initial 1 fps and later 0.2 fps during the slower withdrawal phase. For analysis, integral GFP intensity of image selections limited to cuts were determined in FIJI and plotted as fold change over nuclear background (y-axis) over time (x-axis) to reveal on-off kinetics of FUS (Fig. 2D, F, 4D-G, S3A).

### Treatments and inhibitors

Glycolic acid (GA, Sigma-Aldrich Cat. # 124737) and D-lactate (DL, Sigma-Aldrich Cat. # 71716) were each dissolved in pure, sterile water to obtain 1M stocks, respectively. For GA, 6M NaOH was added drop-wise to assist the dissolution. Both stocks were finally sterile-filtrated. For MNs in MFCs, GA and DL were added together 24 h prior to imaging to both the distal and proximal site at final 1mM each (Fig. 1A, 5A), unless otherwise stated in the titration experiments (Fig. S2A). For laser irradiation experiments and revealing FUS aggregates, GA and DL were added together 24 h prior to imaging to uncompartmentalized dishes at final 10mM each (Fig. 2A, C, E), unless otherwise stated in the titration experiments (Fig. S2A, S3A). For single compound treatments, GA and DL were each added alone to final 20 mM either to both sites in MFCs or to uncompartmentalized dishes (Fig. 1A, 2A, C, S1E, S2A).

Carbonylcyanid-3-chlorphenylhydrazon (CCCP, Sigma-Aldrich Cat. # C2759) and Oligomycin A (Oligo A, Sigma-Aldrich Cat. # 75351) were dissolved in DMSO to obtain a 10 mM stock, respectively. Final working concentrations were 10 µM for each inhibitor. Each inhibitor was added to uncompartmentalized MNs (Fig. 3B) or exclusively to the distal site of MFCs (Fig. 3A) just 4 h prior to imaging to avoid toxic side effects and micro flow progression to the proximal MFC site.

Riluzole (Sigma-Aldrich Cat. # R116) was dissolved in DMSO to obtain a 10mM stock for a final working concentration of 10µM. Culture supernatants were continuously supplemented at both the distal and proximal MFC site with riluzole over the entire MN maturation of 21 days in MFCs prior to imaging (Fig. S4A).

FK866 (Daporinad, Selleckchem Cat. # S2799) was dissolved in DMSO to obtain a 20mM stock for a final working concentration of up to 20µM. Nicotinamide riboside chloride (NAR, Sigma-Aldrich Cat. # SMB00907) was dissolved in sterile culture medium at 100mM for a final working concentration of up to 10mM. Either FK866 or NAR was added to uncompartmentalized cells 24 h prior to imaging (Fig. 4A, C).

DMSO was used as Mock control for CCCP, Oligo A at final 0.1%. Sterile water was used as Mock control for GA and DL.

## Institutional Review Board Statement

The performed procedures were in accordance with the Declaration of Helsinki (WMA, 1964) and approved by the Ethical Committee of the Technische Universität Dresden, Germany (EK 393122012 and EK 45022009)

## Informed Consent Statement

Written informed consent was obtained from all participants including for publication of any research results.

## Author contributions according to CRediT

Conceptualization (AP, AH); Data curation (AP, DG, HG, VZ, RG, ES); Formal Analysis (AP, DG, HG, VZ); Funding acquisition (AH); Investigation (AP, DG, HG, VZ, RG, ES); Methodology (AP, DG, SG, AH); Project administration (AH); Resources (FW, SP, TMB, ES, SG, FP, AH); Software (AP, HG); Supervision (AH); Validation (AP, DG, HG); Visualization (AP); Writing – original draft (AH); Writing – review & editing (all authors)

## Acknowledgments

We acknowledge the great cell culture help of Sylvia Kanzler, Anett Böhme, Katja Zoschke. The Light Microscopy Facility (LMF) of CMCB (Center for Molecular and Cellular Bioengineering, Technische Universität Dresden) provided excellent support for all live imaging experiments. We thank Ronny Sczech for having programmed the original FIJI/KNIME analytical HC organelle trafficking pipeline.

## Movie captions

General information for movie1, 2, 5, 6, 9-12: all movies were acquired with 3 frames per second and channel over 2min (i.e. 400 frames per channel in total). Spinal MNs were cultured in MFCs and stained live with Mitotracker Deep Red or JC-1 (red and green channel simultaneously) and Lysotracker Red to visualize motility of mitochondria and lysosomes in axons. Mitotracker Deep Red is shown in the FIJI look up table (LUT) yellow hot, JC-1 as red/green overlay, Lysotracker Red in cyan hot.

General information for movie3, 4, 7, 8: all movies were acquired with 1 frame per second over 15min. Nuclear UV laser cuts were administered in the BAC HeLa cell model stably expressing normal FUS-eGFP WT or mutant FUS-eGFP P525L or in uncompartmentalized spinal FUS-eGFP Ctrl (wild type) vs isogenic P525L MNs (Table 1) and the recruitment- withdrawal of FUS-eGFP at cuts recorded.

Movie1

Refers to Fig. 1A, showing Mitotracker JC-1 distal (left) vs proximal (right) in FUS as overlay of the red and green channel. Red indicates physiological and green deficient inner membrane potential.

Movie2

Refers to Fig. 1A, showing Lysotracker Red distal (left) vs proximal (right) in FUS.

Movie3

Refers to Fig. 2C, showing recruitment of FUS-GFP to nuclear UV laser cuts in transgenic BAC HeLa cells stably expressing FUS-eGFP Ctrl (wild type) vs P525L.

Movie4

Refers to Fig. 2E, showing recruitment of FUS-eGFP to nuclear UV Laser cuts in spinal FUS-eGFP Ctrl (wild type) vs isogenic P525L MNs.

Movie5

Refers to Fig. 3A, showing Mitotracker JC-1 distal (left) vs proximal (right) in Ctrl spinal MNs under distal Oligo A or CCCP treatment as overlay of the red and green channel. Red indicates physiological and green deficient inner membrane potential.

Movie6

Refers to Fig. 3A, showing Lysotracker Red distal (left) vs proximal (right) in Ctrl spinal MNs under distal Oligo A or CCCP treatment.

Movie7

Refers to Fig. 3B, showing recruitment of FUS-GFP to nuclear UV Laser cuts in spinal FUS- eGFP Ctrl (wild type) MNs treated with Oligo A or CCCP.

Movie8

Refers to Fig. 4C, showing showing recruitment of FUS-eGFP to nuclear UV Laser cuts in transgenic BAC HeLa cells stably expressing FUS-eGFP Ctrl (wild type) vs P525L under NAR or FK866 treatment.

Movie9

Refers to Fig. 5A, showing Mitotracker JC-1 distal (left) vs proximal (right) in SOD1 and TDP43 as overlay of the red and green channel. Red indicates physiological and green deficient inner membrane potential, yellow mixing color a moderate decrease.

Movie10

Refers to Fig. 5A, showing Lysotracker Red distal (left) vs proximal (right) in SOD1 and TDP43.

Movie11

Refers to Fig. S4A, showing Mitotracker Deep Red distal (left) vs proximal (right) in FUS under Riluzole treatment.

Movie12

Refers to Fig. S4A, showing Lysotracker Red distal (left) vs proximal (right) in FUS and SOD1 under Riluzole treatment.

---

**Figure S1.**
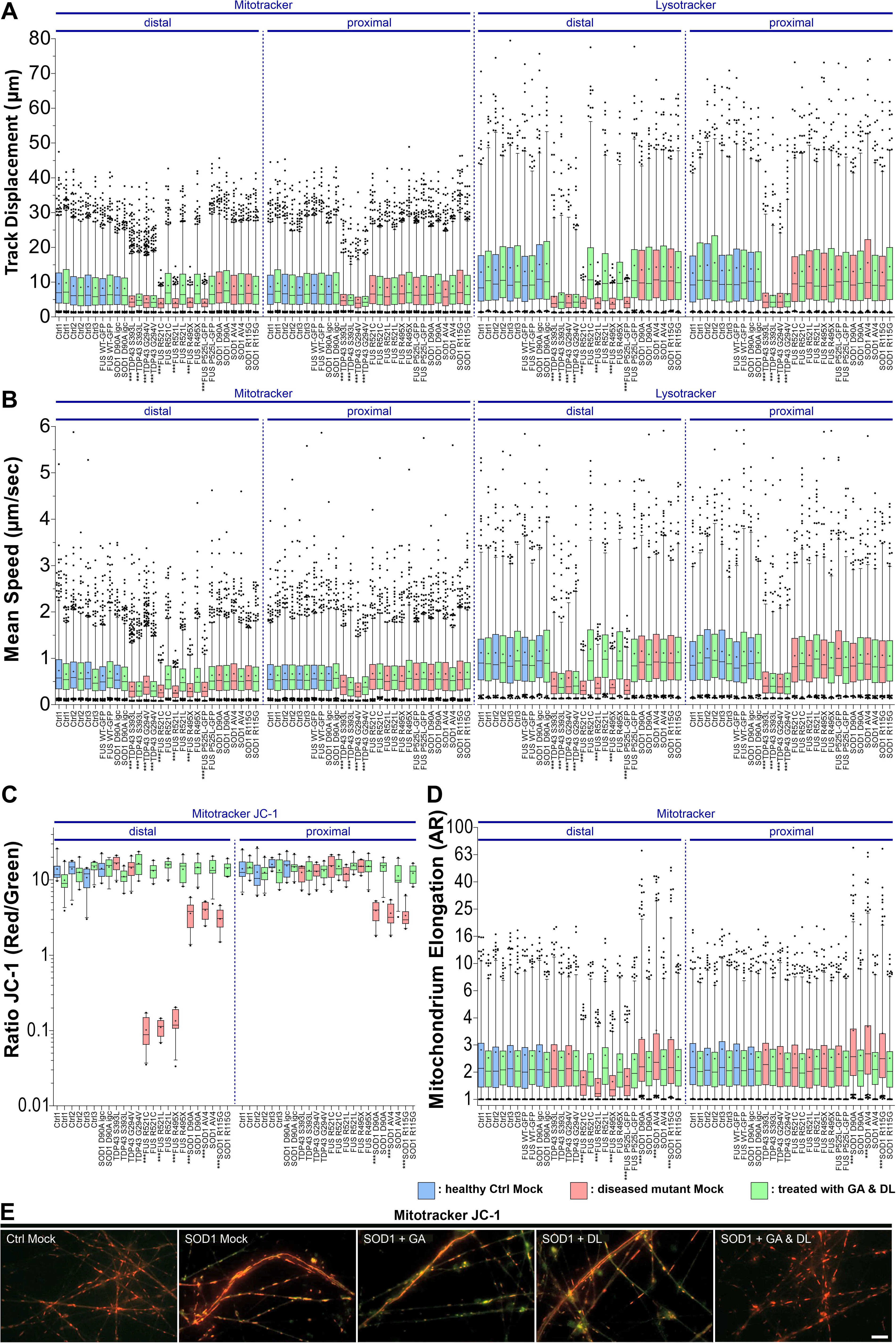
GA and DL together rescue axonal trafficking defects in some but not all ALS mutants. **(A-D)** Refers to Fig. 1 and 5, showing box plot quantifications of all individual mutant (FUS, SOD1, TDP43) and Ctrl lines. A one-way ANOVA with Bonferroni post-hoc test was utilized to reveal significant differences in pairwise comparisons. Asterisks: highly significant alteration compared to Ctrl1 Mock distal (in pale blue), *** p ≤ 0.001, all other pairwise comparisons were not significantly different. **(E)** Super-elongated mitochondria in SOD1 D90A in uncompartmentalized cultures with moderate decrease of inner membrane potential (JC-1 yellow overlap) as compared to Ctrl1. Note the rescue of both phenotypes through GA and DL double (1 mM each) but not GA or DL single treatment (20 mM). Scale bar = 10 µm.

**Figure S2.**
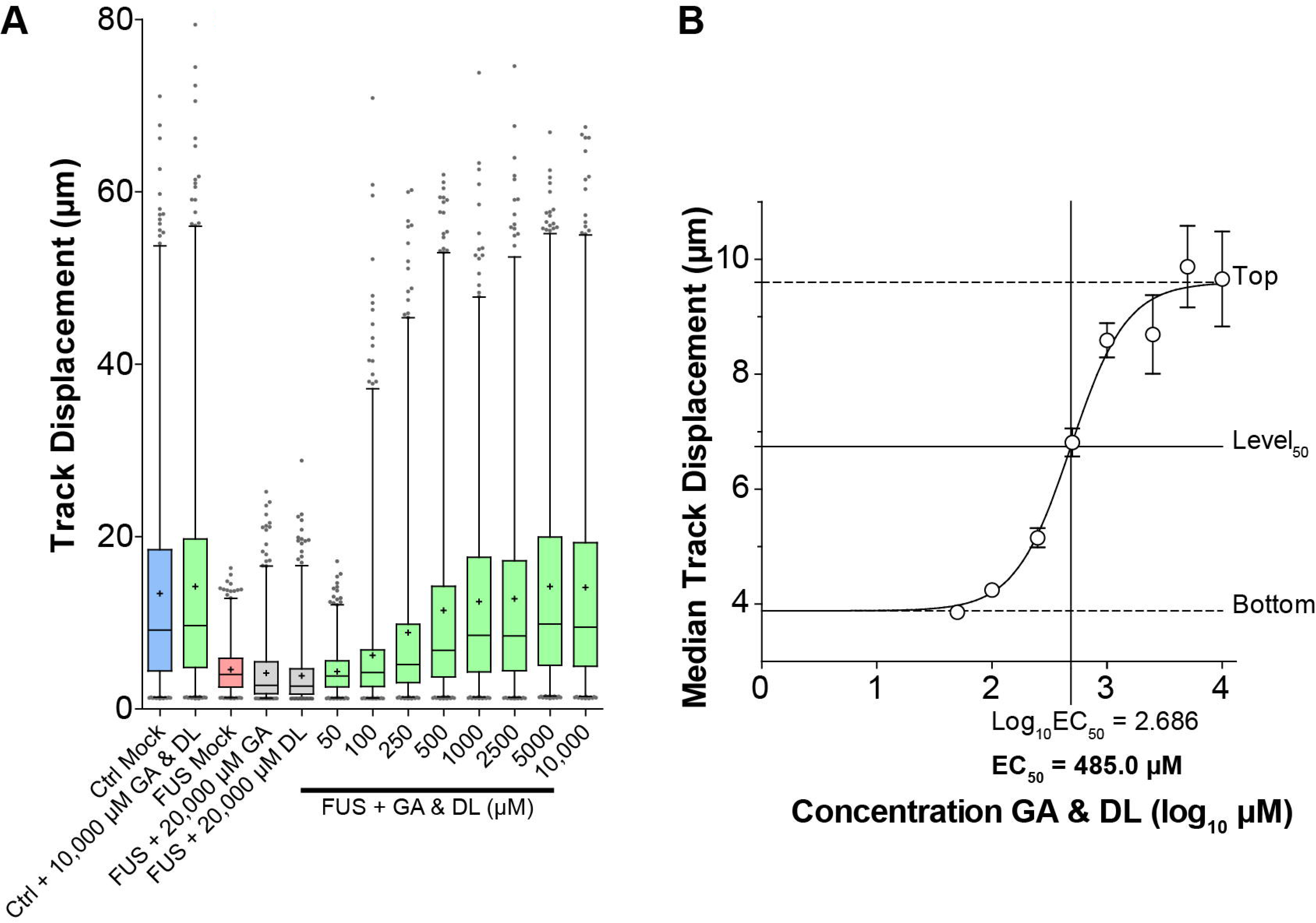
Titration of GA and DL double treatment in rescuing lysosomal trafficking defects in FUS ALS axons reveals an EC_50_ of 485.0 µM. Refers to Fig. 1. Patient-derived isogenic spinal MNs expressing normal (Ctrl) or mutant P525L (FUS) FUS-eGFP (Table 1) were matured in MFCs for 21 DIV. Then the mutant was double-treated for 24 h with GA and DL over a concentration range of 0-10,000µM (10mM) as indicated and imaged live at the distal channel end with Lysotracker. **(A)** Box plots quantifications of track displacement deduced from movies as per organelle values. Box: 25-75% interquartile range, horizontal line: median, cross: mean, whiskers: non-outlier range (99% of data), dots outside whiskers: outliers, Ctrl Mock is shown in pale blue, diseased mutant mock in pale red, double treatment with GA and DL in pale green, single treatments in grey. Note the drastic reduction in distal track displacement in FUS Mock that was rescued through GA and DL double treatment in a concentration-dependent manner with an apparent plateau reached at 1000µM (1mM). Conversely, GA or DL alone did not lead to any increase of organelle track displacement even at 20,000 µM (20 mM). Moreover, double treatment of Ctrl cells did not further stimulate the physiological level even at 10,000 µM (10 mM). **(B)** Non linear regression of (A). Median values of data sets in (A) were plotted over logarithmized GA and DL concentrations to deduce the EC_50_ (x-axis) at level_50_ (y-axis).

**Figure S3.**
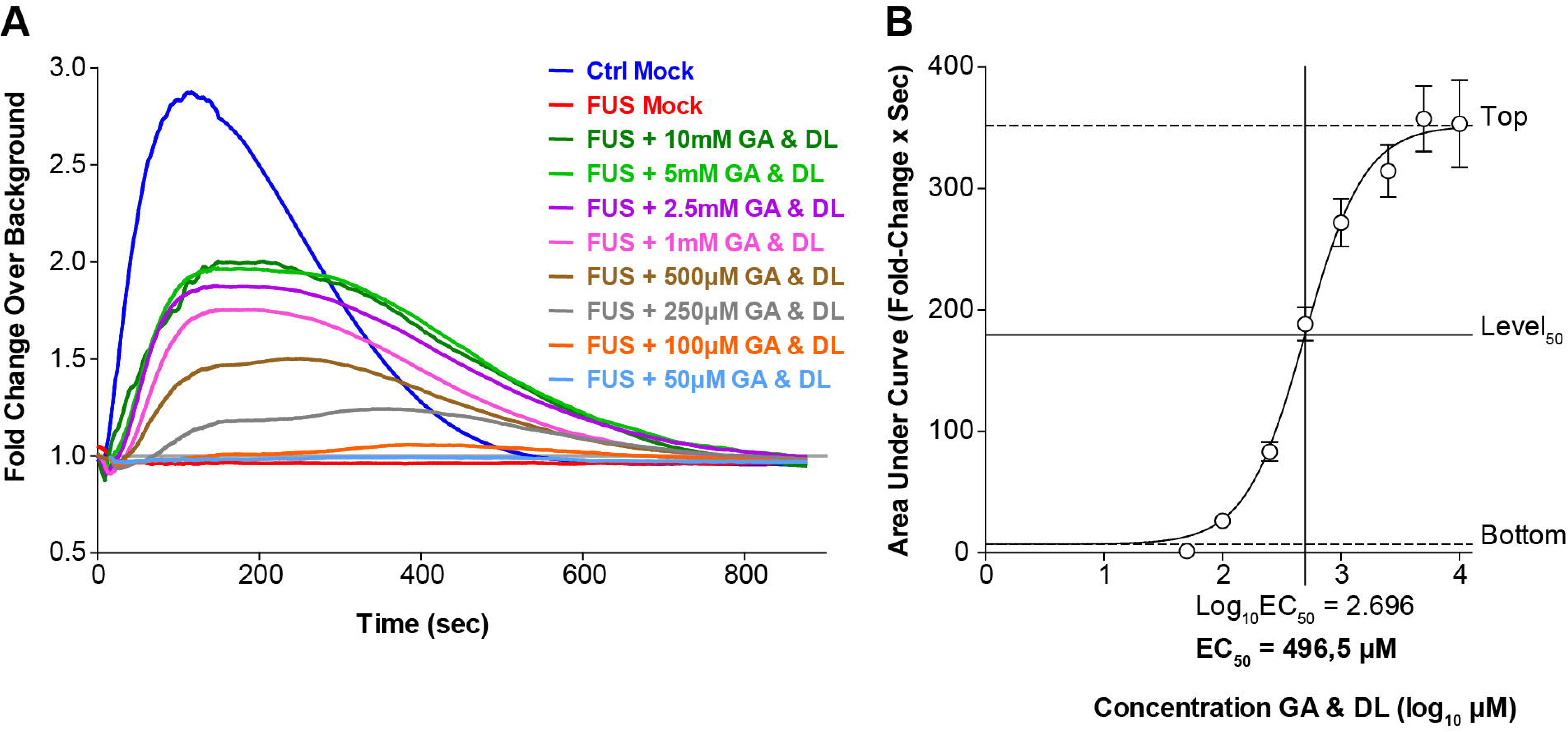
Titration of GA and DL double treatment in rescuing nuclear DNA damage response in FUS ALS reveals an EC_50_ of 496.5 µM. Refers to Fig. 2. Patient-derived isogenic spinal MNs expressing normal (Ctrl) or mutant P525L (FUS) FUS-eGFP (Table 1) were matured for 21 DIV. Then the mutant was double treated for 24 h with GA and DL over a concentration range of 0-10,000 µM (10 mM) as indicated. **(A)** Recruitment-withdrawal of FUS-eGFP to UV Laser cuts in nuclei was then imaged live and the quantified amount of FUS-eGFP at cuts plotted over time. Note the failed recruitment in FUS Mock (red curve) over the entire recording time of 850 sec whereas GA and DL double treatments fairly restored the recruitment-withdrawal towards Ctrl kinetics (blue curve) in a concentration-dependent manner. **(B)** Non linear regression of (A). Areas under curves in (A) were plotted over logarithmized GA and DL concentrations to deduce the EC_50_ (x-axis) at level_50_ (y-axis).

**Figure S4.**
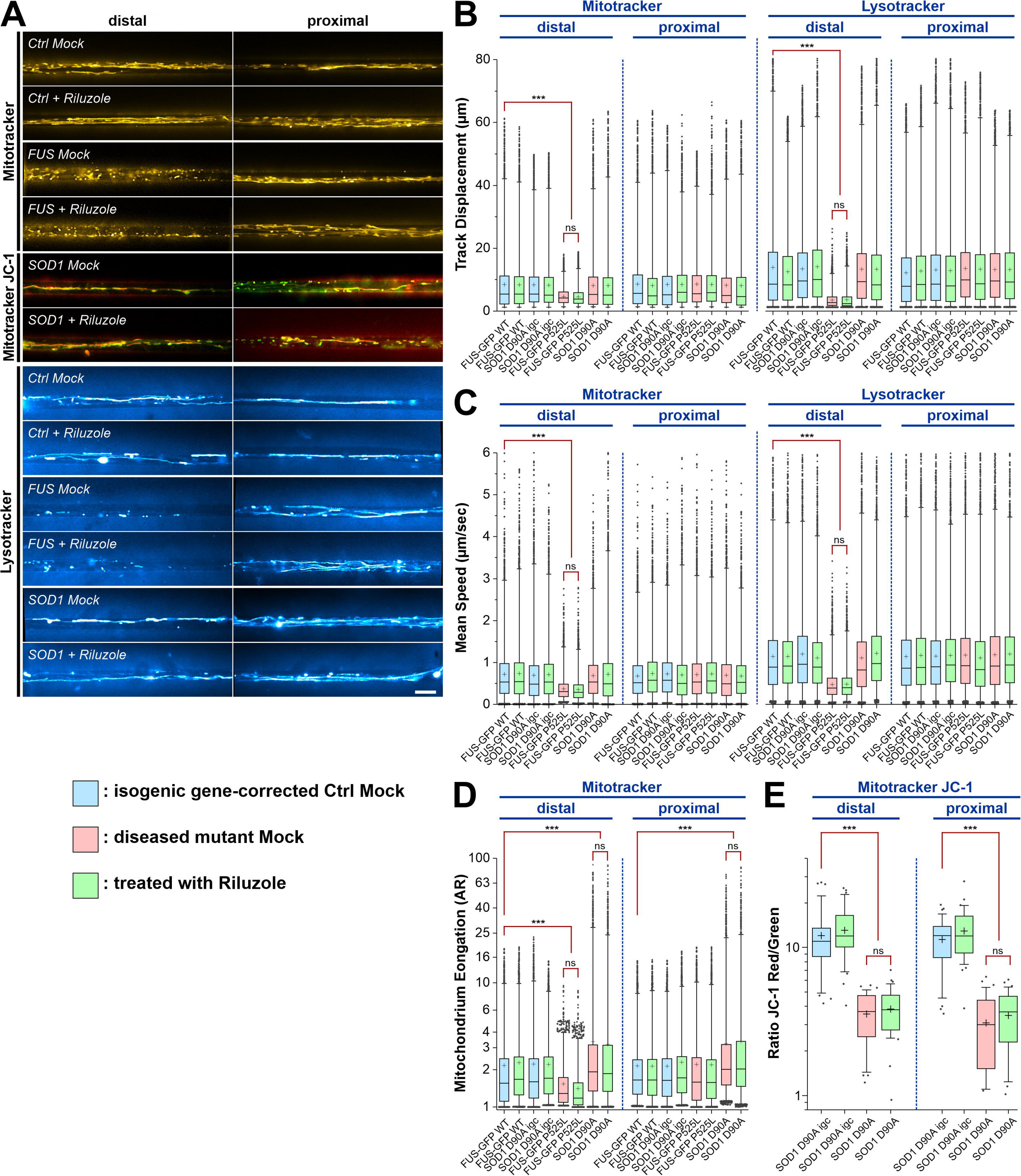
Riluzole fails to restore axonal organelle trafficking in mutant ALS-FUS. **(A)** Maximum intensity projections with Mitotracker and Lysotracker to illustrate axonal organelle motility in mutant FUS (FUS-eGFP P525L, Table 1), SOD1 (SOD1 D90A, Tabel 1) and Ctrl (Ctrl1, Table 1) as in Fig. 1A, 5A (supplementary movies 11 and 12). Note that chronic treatment with Riluzole (Material and Methods) had no effect on neither the distal FUS trafficking defect for both types of organelles nor the abnormal distal and proximal mitochondria elongation and reduced inner membrane potential (JC-1, yellow overlap) in SOD1. Scale bar = 10 µm. **(B-E)** Box plots quantifications of various tracking and morphology parameters deduced from movies from (A) as per organelle values for the mutant SOD1 D90A and FUS-eGFP P525L lines and their isogenic gene-corrected control counterparts SOD1 D90A igc and FUS-eGFP WT (Table 1), except of (E) showing mean values per image. Box: 25-75% interquartile range, horizontal line: median, cross: mean, whiskers: non-outlier range (99% of data), dots outside whiskers: outliers, Ctrl Mock is shown in pale blue, diseased mutant mock in pale red, treatment with Riluzole in pale green. A one-way ANOVA with Bonferroni post-hoc test was utilized to reveal significant differences in pairwise comparisons. Asterisks: highly significant alteration compared to isogenic gene-corrected Ctrl Mock distal (in pale blue), ***p ≤ 0.001, ns: no significant difference, all other pairwise comparisons were not significantly different. **(B, C)** Note the drastic reduction in distal track displacement (B) and mean speed (C) in mutant FUS Mock that was not rescued through Riluzole treatment. **(D)** Note the drastic elongation of mitochondria at both the distal and proximal site in mutant SOD1 that was not rescued through Riluzole treatment. **(E)** Note the reduction of mitochondrial inner membrane potential in mutant SOD1 at both the distal and proximal site that was not rescued through Riluzole treatment.

